# Placentas from SARS-CoV-2 infection during pregnancy exhibit foci of oxidative stress and DNA damage

**DOI:** 10.1101/2024.09.25.614948

**Authors:** Guilherme M Nobrega, Eliza R McColl, Arthur Antolini-Tavares, Renato T Souza, José Guilherme Cecatti, Maria Laura Costa, Indira U Mysorekar

**Affiliations:** Department of Medicine, Section of Infectious Diseases, Baylor College of Medicine, Houston, TX 77030, USA; Department of Obstetrics and Gynecology, School of Medical Sciences, University of Campinas (UNICAMP), Campinas, São Paulo, Brazil; Department of Pathology, School of Medical Sciences, University of Campinas, Campinas, SP, Brazil; Department of Molecular Virology and Microbiology, Baylor College of Medicine, Houston, TX 77030, USA; Huffington Center on Aging, Baylor College of Medicine, Houston, TX 77030, USA

**Keywords:** Cellular Senescence, Cell damage, COVID-19, DNA damage, Reactive Oxygen Species, SARS-CoV-2

## Abstract

**Problem:** COVID-19 during pregnancy is linked to increased maternal morbidity and a higher incidence of preterm births, yet the underlying mechanisms remain unclear. Cellular senescence, characterized by the irreversible cessation of cell division, is a critical process in placental function and its dysregulation has been implicated in pregnancy complications like preterm birth. Senescence can be induced by various stressors, including oxidative stress, DNA damage, and viral infections.

**Method of Study:** In this study, we determined whether COVID-19 had an impact on placental senescence. We examined placentas from women infected with SARS-CoV-2 (n=10 term, 4 preterm) compared to uninfected controls (n=10 term, 3 preterm). The placentas were analyzed for SARS-CoV-2 infection/replication (Spike and Nucleocapsid viral proteins), markers of DNA damage (γH2AX) and oxidative stress (ROS), and senescence (telomere length; cell cycle regulators, SASP).

**Results:** While no overall differences in cellular senescence markers were observed between the COVID-19 positive and negative groups, we found increased secreted SASP markers and confocal microscopy revealed localized areas of oxidative stress and DNA damage in the placentas from COVID-19 positive cases.

**Conclusions:** These findings indicate that SARS-CoV-2 infection induces localized placental damage, warranting further investigation into its impact on maternal and perinatal outcomes.

## INTRODUCTION

The impact of *Severe acute respiratory syndrome coronavirus 2* (SARS-CoV-2) infection among pregnant and postpartum women has been a concern since the beginning of the COVID-19 pandemic.^1^ Severe maternal COVID-19 is associated with an increased risk for preterm birth (PTB) and stillbirth.^2,3^ Although evidence suggests a low risk of vertical transmission and generally mild disease in newborns, the impact on placental health remains poorly understood.^4,5^

The phenomenon of cellular senescence is characterized by cessation of cell growth in response to various stimuli including telomere shortening, oxidative stress, inflammation and cytokines.^6–8^ Senescence plays a critical role in normal placental implantation and development, as well as the onset childbirth. However, dysregulation of senescence has been implicated in obstetric complications such as preeclampsia, preterm labor, PTB, and stillbirth.^6,7,9,10^ In non-placental tissues, SARS-CoV-2 infection has been found to induce senescence, oxidative stress and inflammation.^11^ However, whether SARS-CoV-2 similarly induces senescence in the placenta is unknown. Given its association with obstetric complications, induction of placental senescence by SARS-CoV-2 could impact maternal and perinatal outcomes. Thus, our objective was to determine whether COVID-19 in the placenta is associated with altered markers of senescence characterized by increased expression of the cell cycle inhibitors p21, p53 and p16, DNA damage, activity of the senescence-associated enzyme β-galactosidase (SA-β-gal), and senescence-associated secretory phenotype (SASP) characterized by secretion of inflammatory cytokines/chemokines. To do this, we analyzed markers of senescence in placental and serum samples from unvaccinated pregnant women who were either infected or non-infected with SARS-CoV-2 during pregnancy.

## METHODS

### Ethical considerations

All guidelines for management and use of human samples were followed and approved by the Research Ethics Committee of the University of Campinas, IRB approval number #31591720.5.0000.5404. The participants provided informed consent to sampling, storage and use of clinical samples.

### Placental and serum samples

The present research is a secondary analysis of a large prospective cohort collected by the Brazilian Network of COVID-19 in Obstetrics (REBRACO) from May 2020 to June 2021.^1,12,13^ The present analysis included villous tissue from 27 pregnancies: 10 COVID(+) and 10 COVID(-) cases who delivered at term (TERM), and 4 COVID-19 (+) and 3 COVID-19 (-) cases who delivered preterm (preterm birth, PTB). Exclusion criteria included SARS-CoV-2 vaccination, twin pregnancies, and preeclampsia. Placentas were sampled in an appropriate systematic protocol, defining each sampling site.^14^ Corresponding maternal peripheral blood and umbilical cord blood serum samples (both collected at childbirth) were also obtained.

### DNA and RNA extraction from placental tissue

RNA and DNA were extracted from placental tissue using TRIzol (Invitrogen) according to manufacturer’s protocol and quantified via NanoDrop (Thermo).

### Quantitative reverse-transcription polymerase chain reaction (RT-qPCR)

cDNA was prepared from extracted RNA using SuperScript™ III Reverse Transcriptase (Thermo). mRNA expression of p16, p21 and p53 was quantified using qPCR with SYBR Green (Bio-Rad). YWHAZ was used as an endogenous control. Primer sequences are listed in **Supplemental Data 1**. Relative mRNA expression was obtained using the ΔΔCT method and presented as normalized log_2_ fold change.

### Absolute Telomere Length assay

Telomere length of placental DNA was measured using an Absolute Telomere Length (aTL) assay based on qPCR technique (using primers from IDT and SYBR Green from Bio-Rad), as described previously.^15^

### Luminex analysis of maternal and umbilical cord serum

Levels of the SASP markers VEGFA, IL-1α, MIP-1α, IL-6, IL-8, and GRO-α were evaluated in maternal serum and umbilical cord serum by Luminex assay (Procartaplex – Thermo). Luminex assay was performed at BCM Proteomics Core according to manufacturer specifications.

### Immunofluorescence in Placental Tissue

Paraffin-embedded placental tissue was cut into 5 µm thick sections. The sections were deparaffinized using Histoclear and hydrated in an alcoholic series until reaching distilled water (ddH₂O). After hydration, antigen retrieval was performed using a 10 mM citrate buffer at pH 6.0 with 0.05% Tween-20 solution in a pressure cooker for 1 hour (warming up, 5 minutes under pressure, and depressurizing). After cooling down, blocking was carried out using 5% horse serum in 1X PBS with 0.3% Triton-X. Sections were incubated overnight at 4℃ with primary antibodies specific for SARS-CoV-2 Nucleocapsid (1:300, Invitrogen), Spike (1:500, Invitrogen), 8-oxoguanine (1:50, Abcam) and γH2AX (1:300, Cell Signaling) in a solution of 5% Horse serum in 1X PBS with 0.2% Triton-X. Following washing in PBS, sections were stained with anti-mouse and anti-rabbit Alexa Fluor-conjugated secondary antibodies (1:500-1:1000, Invitrogen) for one hour at room temperature. Autofluorescence was quenched using VectaShield antifading kit (VectorLabs) prior to counterstaining with Hoescht nuclear stain (1:10,000). Slides were visualized using an Eclipse Ti2 Nikon confocal microscope at 20 and 40x with Galvano capture. For quantification, the mean intensity of fluorescence of 5 random fields at 20x were averaged for each slide.

### Protein extraction and Western Blotting

Protein lysates were extracted from 75 mg of placental tissue with T-PER (Thermo) buffer and quantified by BCA (Pierce™ BCA Protein Assay Kit, Thermo) Proteins were separated by 12% SDS-PAGE and transferred to PVDF membranes. After blocking with Intercept® TBS Protein-Free Blocking Buffer (LICORbio), membranes were incubated with anti-γH2AX (1:500, Cell Signaling) or anti-GAPDH (1:1000, Cell Signaling) primary antibodies overnight at 4℃ followed by anti-rabbit IRDye 800CW secondary antibody (1:3000, Invitrogen) for one hour at room temperature. Bands were visualized with a ChemiDoc (BioRad) and intensity was determined using ImageLab 6.1 (BioRad). Intensity of γH2AX was normalized to that of GAPDH.

### Data analysis

Comparisons between groups were performed using Mann-Whitney test (U test), with *p* ≤ 0.05 considered significant. Outliers were considered by the ROUT method Q = 0.2%. All statistical tests were performed using GraphPad Prism version 7 for Mac (GraphPad Software).

## RESULTS AND DISCUSSION

Within the infected cohort, SARS-CoV-2 Nucleocapsid and Spike protein were detected in the villous tissue of term and PTB placentas of COVID-19 (+) pregnancies (**Figure 1A**). Nucleocapsid (green) and Spike (red) proteins colocalized in syncytiotrophoblasts, with additional puncta of Spike protein in the villous stroma, consistent with what was shown previously.^16^ We found no significant difference in mRNA expression of p16, p21, p53 between COVID-19 (+) and (-) cases in either term or PTB placentas (**Figure 1 B-D**). The p16, p21, p53 heat map exhibit this pattern according to a scale considering the gestational age at childbirth, presenting the similar pattern of infected and uninfected cases (**Figure 1 B-E).** However, p21 mRNA is significantly decreased in PTB placentas (*p-value* = 0.0160), regardless of COVID-19 status, indicative of increased senescence in PTB placentas as previously described.^17^ Preterm placentas had shorter telomeres compared to term placentas as noted previously ^18^ and was not affected by COVID-19 status (p-value < 0.0001) (**Figure 1 F**). Luminex assay for detecting changes in SASP in sera revealed that GRO-α was significantly increased in maternal serum (**Figure 1 G**) and VEGFA was significantly increased in umbilical cord serum (**Figure 1 H**) in COVID-19 (+) cases. However, we found no significant difference in levels of SASP markers in maternal and umbilical cord serum between COVID-19 (+) and (-) cases (**Supplemental Figure 1**).

**Figure 1.**
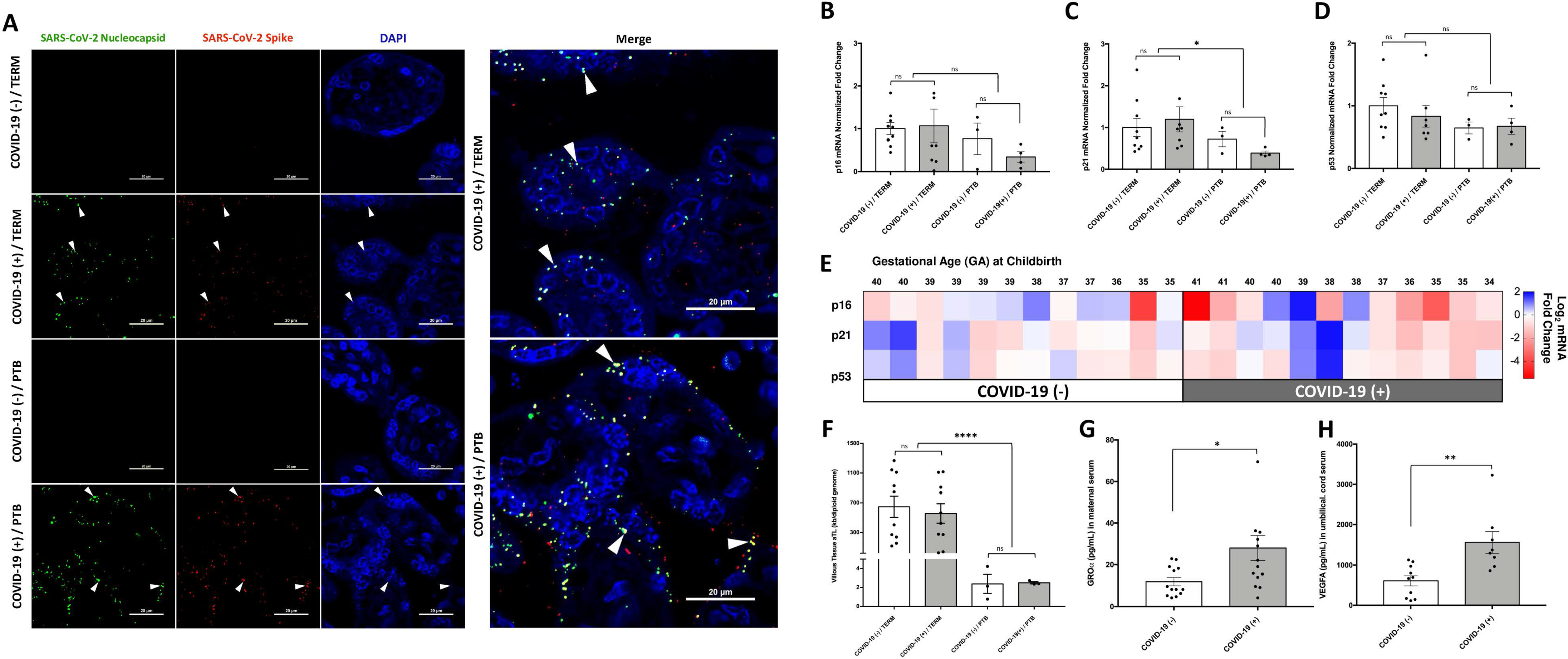
SARS-CoV-2 detection and senescence pattern in SARS-CoV-2 positive placentas. **(A)** SARS-CoV-2 nucleocapsid (green) and spike (red) proteins colocalization (yellow in merge) in villous tissue of term (TERM) and preterm (PTB) COVID-19 (+) pregnancies – 40 X magnification focus, DAPI (blue). **(B-E)** p16 **(B)**, p21 **(C)** and p53 **(D)** relative mRNA expression in placental tissue of PTB/TERM COVID-19(+)/(-) pregnancies; bars represent means and error bars represent standard error of the mean (SEM); *: *p-value* < 0.05; ns: non-significant difference. **(E)** Heat map of p16, p21 and p53 relative mRNA expression adjusted by gestational age at childbirth in completed weeks in COVID-19 (-)/(+) pregnancies. **(F)** Absolute Telomere Length (aTL) in placental tissue of PTB/TERM COVID-19(+)/(-) pregnancies; bars represent means and error bars represent standard error of the mean (SEM); ***: *p-value* < 0.0001; ns: non-significant difference. **(G-H)** GRO-α in maternal serum **(G)** and VEGFA in umbilical cord serum **(H)** considering COVID-19(+)/(-) pregnancies; bars represent means and error bars represent standard error of the mean (SEM). *: *p-value* = 0.0199; **: *p-value* = 0.0014.

DNA damage was assessed by ɣH2AX) and oxidative stress (assessed by 8-oxoguanine, a product of DNA-ROS interaction). COVID-19 was not associated with significant changes in quantification of ɣH2AX (**Figure 2 B, C**) and 8-oxoguanine (**Figure 2 E**) overall. However, we observed highly localized foci of oxidative stress and DNA damage in SARS-CoV-2 infected placentas with punctate staining of ɣH2AX (**Figure 2 A**) and 8-oxoguanine (**Figure 2 D**) observed in both term and PTB cases. While COVID-19 (-) placentas exhibited nuclear localization of ɣH2AX and limited 8-oxoguanine, COVID-19 (+) cases had foci in the STBs in which 8-oxog puncta colocalized with extra-nuclear ɣH2AX (**Figure 2 F**). This pattern recapitulates findings in pathological analysis of SARS-CoV-2 infected women placentas characterized by focal changes.^19,20^

**Figure 2.**
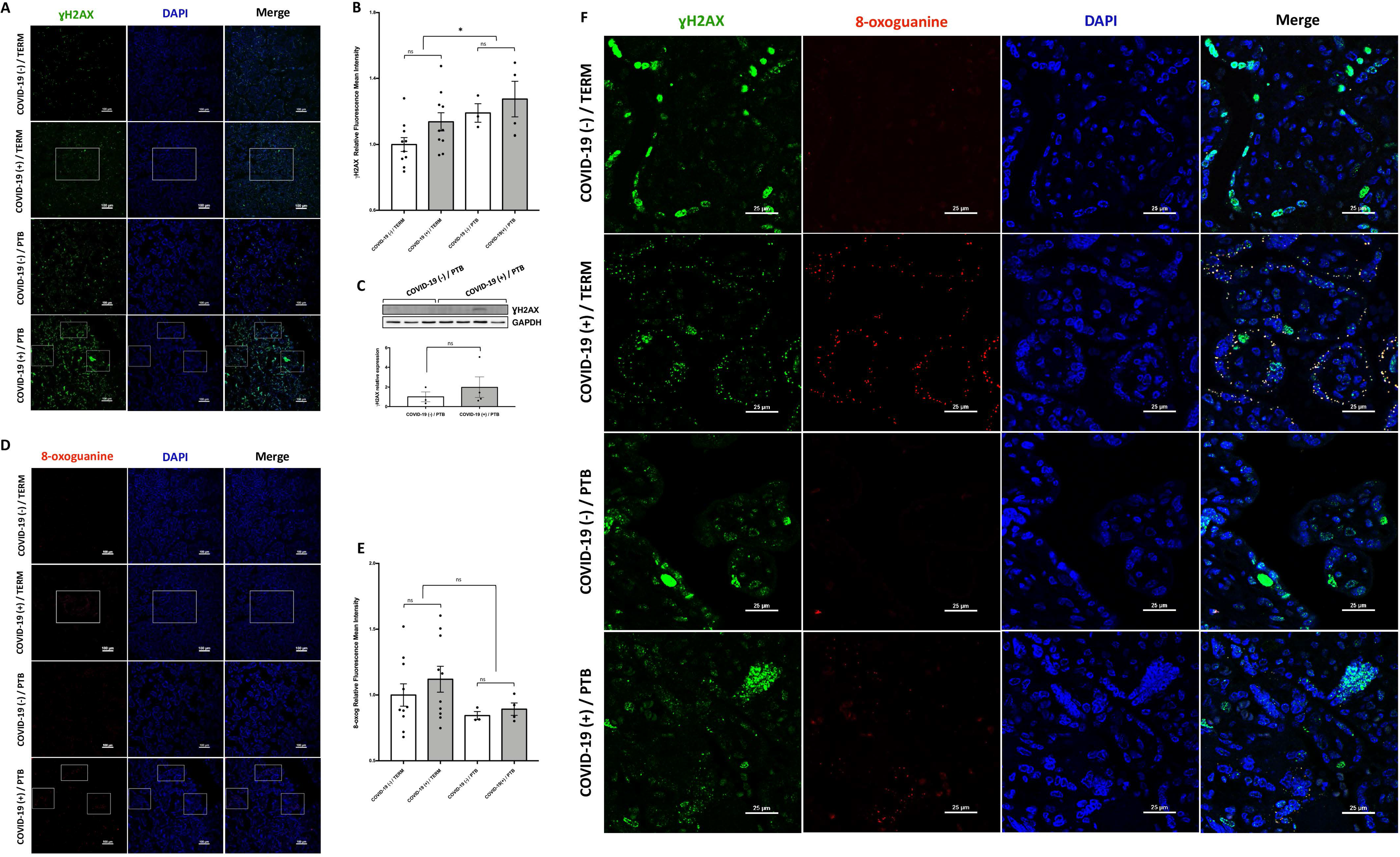
DNA damage and ROS activity punctate foci in SARS-CoV-2 positive placentas. **(A) γ**H2AX in villous tissue of term (TERM) and preterm (PTB) COVID-19 (+) pregnancies – 20 X magnification, DAPI (blue). **(B) γ**H2AX relative fluorescence mean intensity in placental tissue of PTB/TERM COVID-19(+)/(-) pregnancies; bars represent means and error bars represent standard error of the mean (SEM); *: *p-value* < 0.05; ns: non-significant difference. **(C) γ**H2AX protein relative quantification in COVID-19 (+/-)/PTB villous tissue; bars represent means and error bars represent standard error of the mean (SEM); ns: non-significant difference **(D)** 8-oxoguanine (red) in villous tissue of TERM and PTB COVID-19 (+) pregnancies – 20 X magnification, DAPI (blue). **(E) γ**H2AX **(C)** and 8-oxoguanine relative fluorescence mean intensity in placental tissue of PTB/TERM COVID-19(+)/(-) pregnancies; bars represent means and error bars represent standard error of the mean (SEM); *: *p-value* < 0.05; ns: non-significant difference. **(F) γ**H2AX (green, DNA damage) and 8-oxoguanine (red, ROS activity) colocalization (yellow in merge) in punctate *foci* in TERM SARS-CoV-2 positive placentas – 40X magnification, DAPI (blue).

In this report, we demonstrate that COVID-19 (+) placentas show a distinct phenotype of foci of oxidative stress and DNA damage. These changes, along with the presence of SARS-CoV-2 viral proteins in the villi that persist in the placenta after infection, may serve as a long-term marker of SARS-CoV-2 damage in placental tissue, analogous to a “scar” induced by previous infection. In addition, few inflammatory secreted senescence markers in umbilical cord blood and maternal blood serum were affected by COVID-19. Given the focal impact of SARS-CoV-2, as also observed in the context of preeclampsia,^21^ studies of the placenta should consider infection heterogeneity, especially when utilizing techniques like single-cell RNA sequencing and other approaches. Organoid models of infection and *in vivo* approaches may provide mechanistical insights necessary to understand the molecular basis of SARS-CoV-2 focal disruption and damage in placental tissue.

There were no overall changes in the quantity of senescence markers in placentas from unvaccinated women infected with SARS-CoV-2. Thus, the observations regarding senescence markers associated with SARS-CoV-2 infection observed in human lung cell lines and explants, as well in human brain cell lines, were not replicated in the same pattern in this subset of human placental samples.^22,23^ The placenta possesses a range of compensatory mechanisms to cope the infection and measuring the impact of senescence in an organ that is already developing its markers in a third trimester context remains a challenge.^24^

Our study has a few limitations. First, the samples were collected at a single center, focusing on a geolocated population primarily from medium to high-density urban areas, all assessed at a tertiary care facility. Second, the cases selected for analysis were subject to exclusion criteria that, while refining the potential findings, limited the sample size due to the high-risk nature of the population evaluated.

## ACKNOWLEDGEMENTS

GMN was supported by Brazilian Coordination of Superior Level Staff Improvement (CAPES) [grant number 88887.712761/2022-00], by CAPES Institutional Internationalization Program (CAPES/PrInt) [grant number 88887.891986/2023-00] and Santander Bank Short-Term International Fellowship [University of Campinas 2023 grant]. MLC was supported by São Paulo Research Foundation (FAPESP) [grant number 2021/09937-1] and by Brazilian National Council for Scientific and Technological Development (CNPq) [grant number 408407/2021-2 and 308378/2022-9]. IUM and MLC were also supported by the Washington University at Saint Louis, USA, McDonnell Academy seed grant for research on infectious diseases and the impact of COVID-19. We thank all the REBRACO Study Group collaborators, as the core part of the current study.

## AUTHOR CONTRIBUTIONS STATEMENT

GMN, ERM and IUM developed the concept of the study. GMN, AA-T, RTS, JGC and MLC implemented clinical data collection and biological sampling and adequate storage. GMN and ERM performed all the laboratorial analyses. GMN and ERM ran the statistical analyses. GMN wrote the first manuscript draft, and all authors further reviewed the manuscript. IUM and MLC supervised all the project analyses. IUM and MLC coordinated the study and were responsible for the project administration and funding acquisition. The collaboration between all authors was fundamental for biological sampling, clinical data collection and work development. All authors have read and agreed to the published version of the manuscript.

## ETHICS STATEMENT

The authors confirm that the ethical policies of the journal, as noted on the journal’s author guidelines page, have been adhered to. All guidelines for management and use of human samples were followed and approved by the Research Ethics Committee of the University of Campinas, IRB approval number #31591720.5.0000.5404. The participants provided informed consent to sampling, storage and use of clinical samples.

---

**Supplemental Figure 1.**
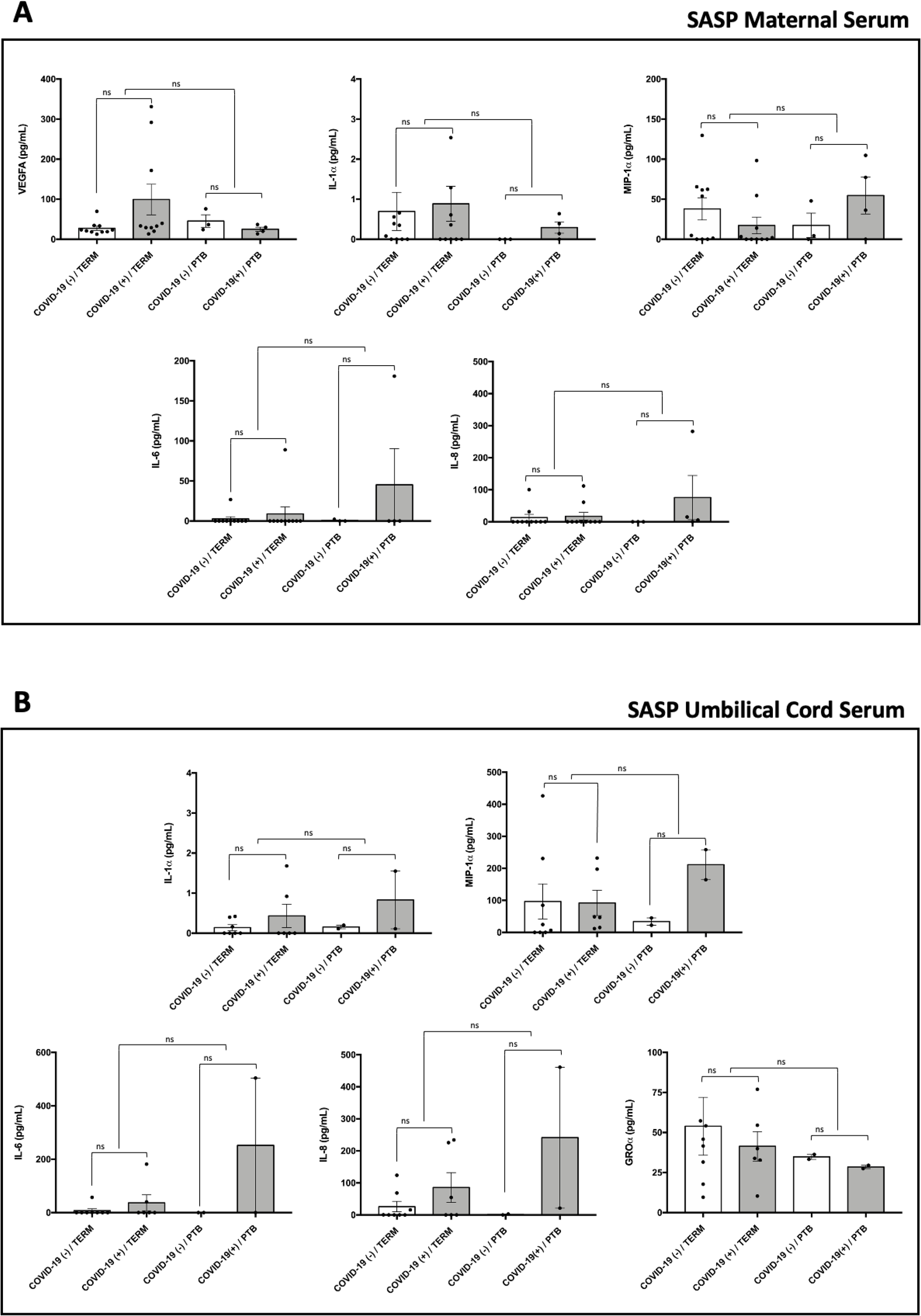
Senescence associated secretory associated (SASP) profile in Maternal Serum **(A)** and Umbilical Cord Serum **(B)** considering VEGFA, IL-1α, MIP-1α, IL-6, IL-8 and GRO-α; bars represent means and error bars represent standard error of the mean (SEM); ns: non-significant difference.

**Supplemental Table 1.**
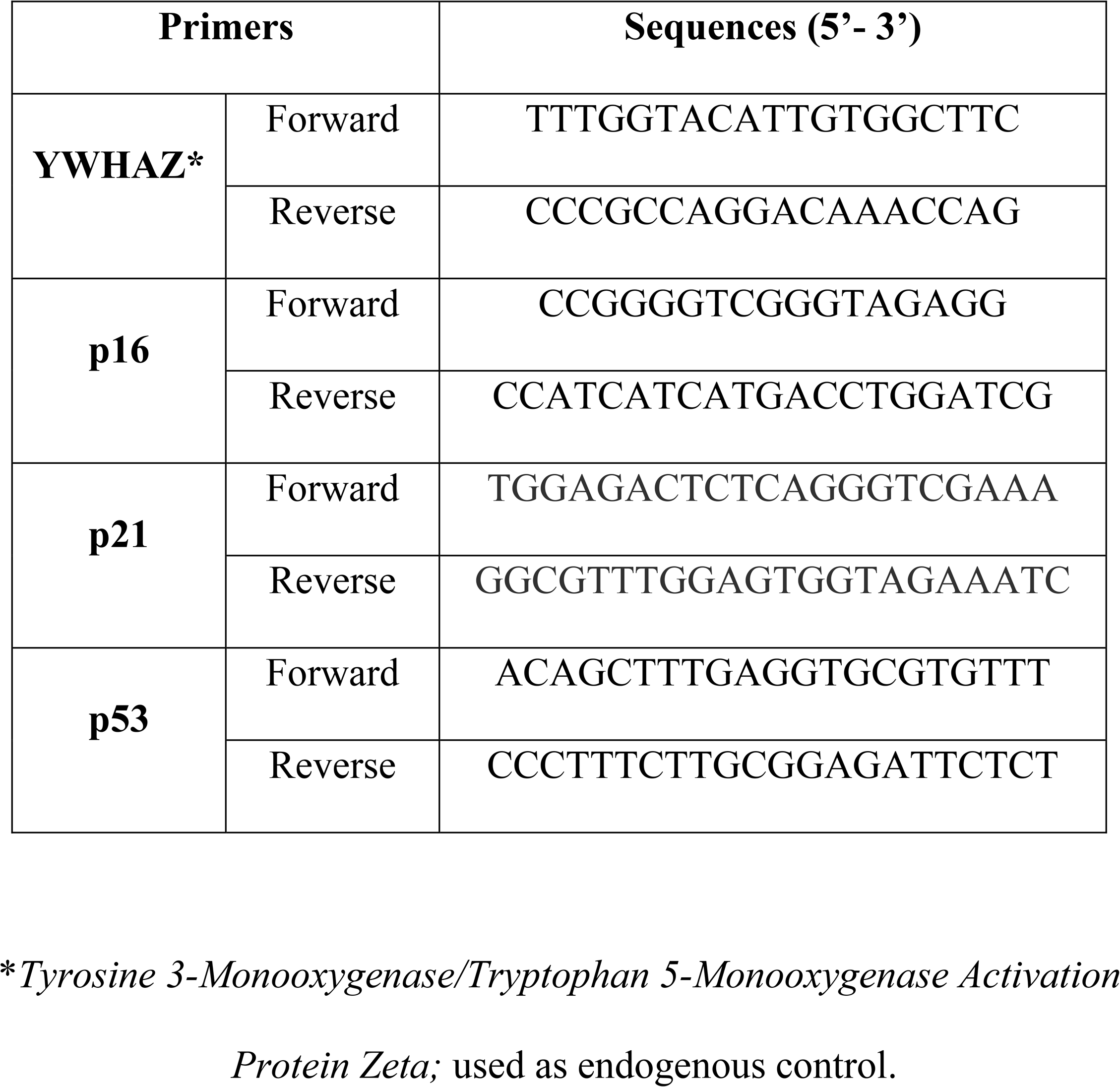
Table of primers sequences and cycling conditions for real-time PCR (qPCR) analysis with SYBR Green detection.

